# Point-Cloud Enhancement and Structural Interpretation Framework for Cryo-EM Maps

**DOI:** 10.64898/2026.01.17.700079

**Authors:** Konrad Karanowski, Miłosz Chojecki, Andrzej Rzepiela, Mateusz Grzesiuk, Radosław Kuczbański, Lukas Frey, Maciej Zięba

## Abstract

Cryo-Electron Microscopy (cryo-EM) has become a cornerstone of modern structural biochemistry, enabling the reconstruction of 3D protein maps at near-atomic resolution. Despite its transformative impact, interpreting maps remains challenging. Structural heterogeneity and molecular flexibility often produce low-resolution densities, which post-processing can only sharpen in well-ordered regions. This leaves flexible areas either poorly resolved or lost. Recent deep-learning approaches have demonstrated strong potential for enhancing cryo-EM maps, enabling an extended interpretation of cryo-EM densities. Most of these methods operate on small volumetric blocks, which restricts the receptive field of the model and prevents it from leveraging the broader structural context of the protein. To address this limitation, we introduce *CryoPC*, a model that enriches local block-based processing with a compact point-cloud representation of the entire map. By conditioning the network on this global representation, CryoPC is able to capture both local and distant structural features, allowing it to make more globally consistent predictions. We demonstrate that incorporating this global context consistently improves enhancement quality across a variety of samples. Furthermore, CryoPC achieves great performance compared to existing methods of similar speed, offering a practical and scalable solution for cryo-EM map enhancement. Finally, we present a framework for assesing a quality of enhanced maps using metrics from the Phenix package.

## Introduction

Cryo-Electron Microscopy (cryo-EM) is an imaging technique that enables the reconstruction of three dimensional protein structures with nearly atomic resolution (Kühlbrandt, 2014). In contrast to other structural methods such as X-ray crystallography, cryo-EM allows the observation of biomolecules in a state close to their native environment, as well as the visualization of multiple conformations like F-ATPase within one sample (Courbon and Rubinstein, 2022). On the other hand for example nuclear magnetic resonance (NMR) is limited in protein size and depends on isotopic labeling strategies to study proteins in solution. Cryo-EM advantages besides small sample volume are the ability to extract native proteins such as the large nuclear pore complex isolated from *Xenopus laevis* oocyte (Fontana et al., 2022), ribosomes or viruses such as rotavirus (Chen et al., 2009) or the spike protein of SARS-CoV-2 (Shi et al., 2023). The workflow leading to a three-dimensional reconstruction (cryo-EM map) typically involves multiple steps: started with protein purification and sample vitrification (Dubochet et al., 1988), followed by high-throughput cryo-EM data acquisition using a transmission electron microscope (TEM) (Frank, 2006). After motion-correction the particles are picked and currated, (Bepler et al., 2019; Dhakal et al., 2024, 2025; Wagner et al., 2019) (Kimanius et al., 2021) followed by initial model generation, 3D refinement, reconstruction and post-processing of the cryo-EM map (Punjani et al., 2017; Zhong et al., 2021). This cryo-EM map can then be used for *de novo* atomic model building (Terashi and Kihara, 2018; Terwilliger et al., 2018a; Jamali et al., 2024). However, the resolution of 3D reconstructions is constrained by multiple sample- and method-dependent factors such as poor signal-to-noise ratio (SNR ~ 0.1), protein mass dependent contrast, and frequency-dependent attenuation by the contrast transfer function (CTF) (Palovcak et al., 2020; Bendory et al., 2019). These limitations create a need for effective post-processing techniques,such as map-sharpening and map-enhancement methods.

Global map sharpening typically applies a single B-factor correction to the entire reconstruction based on Guinier analysis (Rosenthal and Henderson, 2003) in an effort to enhance the clarity of structural features. Because the overall level of detail present in the map directly influences the accuracy of the resulting atomic models, such global approaches are widely used in software packages including `phenix.auto_sharpen` (Terwilliger et al., 2018b), the post-processing module in RELION (Scheres, 2012), and the sharpening utilities available in CryoSPARC (Punjani et al., 2017). A key limitation of these methods is the assumption that a single B-factor appropriately describes the entire map. In practice, cryo-EM reconstructions often display substantial spatial variability in resolution and noise characteristics. As a result, uniform sharpening can leave some regions insufficiently enhanced while pushing others into over-sharpened, noisy regimes, ultimately reducing map interpretability.

Local sharpening methods attempt to overcome this issue by adapting the sharpening strength to the characteristics of individual regions of the density map. For example, LocalDeblur (Ramírez-Aportela et al., 2019) performs Wiener-filter–based deblurring with weights that depend on the locally estimated resolution. LocScale (Jakobi et al., 2017), on the other hand, adjusts local amplitudes using a sliding-window strategy to bring the density map into agreement with an associated atomic model. Although such approaches represent an improvement over global sharpening, they are still limited in their ability to accommodate the full complexity and heterogeneity present in modern cryo-EM maps.

With the rapid progress of deep learning, several neural network–based approaches for cryo-EM map enhancement have been introduced. DeepEMEnhancer (Sánchez-García et al., 2020) employs a UNet architecture (Ronneberger et al., 2015) trained to approximate the effect of LocScale by learning from pairs of experimental maps and their LocScale-processed counterparts. However, because the target maps are themselves derived from the noisy experimental data, the method inherits the limitations of LocScale and cannot fully overcome errors present in the original reconstructions. An alternative strategy is to train models using experimental maps paired with simulated, noise-free maps generated from reference atomic structures using Gaussian-based forward models (DiMaio et al., 2009; Berman, 2000). EMReady (He et al., 2023), for example, leverages a Swin-Conv-UNet network (Zhang et al., 2023) to improve a wide range of cryo-EM maps. CryoTEN (Selvaraj et al., 2025) is using a convolution-transformer inspired by UNETR++ (Shaker et al., 2022) for efficient and robust 3D enhancement. CryoSAMU (Zhang et al., 2025) incorporates cross-attention mechanisms (Vaswani et al., 2023; Petit et al., 2021) and conditions its network on ESM-IF1 embeddings (Hsu et al., 2022) to refine structural features during training. Despite their successes, all of these methods share a fundamental limitation: due to the large and variable size of cryo-EM maps, they operate only on small volumetric patches (typically 48^3^ – 64^3^ voxels). As a consequence, networks capture only local information and lack awareness of the global structural context, which can lead to inconsistencies or missed long-range features.

To address these limitations, we introduce **CryoPC**, a cryo-EM map enhancement method that combines local volumetric learning with global structural context. The method is based on a UNet backbone for efficient multi-scale density refinement, augmented with a global conditioning mechanism that captures long-range structural information. Global context is obtained by converting the full cryo-EM map into a sparse 3D point-cloud representation, which is processed by a PointNet (Qi et al., 2017) module to extract a compact global descriptor. This representation conditions the UNet through a cross-attention mechanism, enabling local density enhancement to be guided by the overall macromolecular topology. Our contributions are three-fold: **(i)** we introduce a novel point-cloud–conditioned cryo-EM map enhancement framework, **(ii)** propose a map–model–based evaluation strategy to assess structural fidelity, and **(iii)** demonstrate improved map quality with fast inference suitable for large-scale cryo-EM workflows.

## Material and methods

### Data collection and processing

We constructed a dataset of cryo-EM map–model pairs following the pipeline and data list provided by Selvaraj et al. (2025). Single-particle cryo-EM structures from 2 to 7 Å resolution range were collected from RCSB Protein Data Bank (PDB). When multiple maps were associated with a single PDB entry, one representative map was retained, and likewise only one model was selected when multiple structures corresponded to the same map. For each selected structure, the corresponding protein PDB file and primary cryo-EM map (post-processed) were downloaded from the PDB or EMDB. Map–model cross-correlation (CC) values were computed using `phenix.map_to_model_cc` (Afonine et al., 2018), and only maps with CC-mask *>* 0.7 and CC-box *>* 0.6 were kept to ensure data quality. To reduce redundancy, we clustered sequences with MM-seqs2 (Steinegger and Söding, 2017) at 30% sequence identity and selected a single representative structure from each cluster. This procedure yielded 1521 non-redundant map–model pairs. From this dataset, 1295 pairs were used for training, 76 for validation, and 120 for testing.

Training data was generated according to (Selvaraj et al., 2025). The deposited experimental cryo-EM maps were used as network inputs, and paired high-quality simulated maps derived from their corresponding PDB structures served as supervision targets. Simulated maps were generated using the Gaussian-based density formulation (DiMaio et al., 2009):

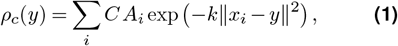

where *x*_*i*_ and *A*_*i*_ denote the coordinates and atomic numbers of atoms in the PDB structure, *k* = *π/*(0.9*R*_0_)^2^, and *R*_0_ is the FSC_model_ at 0.143 resolution computed with `phenix.mtriage` (Afonine et al., 2018). The normalization constant is *C* = (*k/π*)^3*/*2^. Densities were evaluated at all grid points using all atoms in the structure, including hydrogen and nonstandard residues.

Both the experimental and simulated maps were resampled to a uniform voxel spacing of 1Å. Experimental maps were normalized to the range 0-1 using the 99.999th-percentile density value. For the training set, maps were first divided into overlapping 64^3^ cubes with a stride of 50 voxels, and 48^3^ sub-cubes were randomly cropped on the fly to reduce overfitting. Only cubes containing protein density were retained to avoid noise bias. For validation and testing, maps were directly split into overlapping blocks of fixed size 48^3^ using a stride of 38 voxels.

### 3D Point Cloud representation

Point clouds serve as a sparse and memory-efficient representation of 3D structures, enabling the modeling of complex geometries without the computational cost associated with processing dense volumetric grids (Sarker et al., 2024). By focusing only on the occupied space, point clouds allow deep learning models to capture global topological information and long-range dependencies, which are typically lost when analyzing small, localized volumetric patches independent of one another.

To incorporate this global context into our enhancement process, we generate a point cloud representation directly from the raw cryo-EM density map. Our generation pipeline follows a statistical thresholding and sampling strategy. First, given an input volume *V*, we calculate the global mean intensity *μ* and standard deviation *σ*. We define a density threshold *T* as:

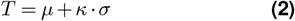

where *κ* is a threshold factor. Voxels with density values below *T* are discarded as background noise, while voxels exceeding *T* are considered potential structural features.

From the set of remaining active voxels, we randomly sample a fixed number of points *N* to ensure a consistent input size for the network. If the number of active voxels exceeds *N*, we perform subsampling without replacement; otherwise, we sample with replacement to meet the required count. Unlike standard geometric point clouds that typically consist only of spatial coordinates (*x, y, z*), our method utilizes a 4-dimensional representation. Each point *p*_*i*_ is defined as a tuple (*x*_*i*_, *y*_*i*_, *z*_*i*_, *v*_*i*_), where *x, y, z* denote the spatial coordinates within the grid, and *v*_*i*_ represents the density intensity at that location. We used 512 points with a threshold factor *κ* of 1.5.

### Our method

CryoPC adopts a UNet-style architecture, structurally outlined in Figure 2. The model operates on two parallel branches: a convolutional encoder that processes local volumetric patches, and a PointNet (Qi et al., 2017) module that extracts a global feature vector from the point cloud representation to modulate the latent space. The encoder begins with a 3D projection convolution layer that processes the single-channel input volume. This is followed by three stages, each containing two residual convolutional blocks and a four-headed linear self-attention layer. Each residual block comprises two sub-blocks, and each sub-block consists of group normalization (Wu and He, 2018), a SiLU activation function (Elfwing et al., 2017), a dropout layer with *p* = 0.2, and a convolution layer. A fourth encoder stage includes only two residual convolutional blocks and serves as the bottleneck representation.

The cross-attention (Vaswani et al., 2023) module is composed of a residual block, followed by a four-headed linear self-attention layer, a cross-attention layer, and a second residual block. To obtain a spatial representation, we employ PointNet (Qi et al., 2017), consisting of a single T-Net followed by three 1D convolutional layers, an argmax aggregation, and a final linear layer. The network accepts 4D point cloud representation (x-coordinate, y-coordinate, z-coordinate and point value) and returns single feature vector. Cross attention is computed between this vector and the encoder representation of a volume.

The decoder mirrors the encoder structure. Feature maps are progressively upsampled using nearest-neighbor interpolation and concatenated with the corresponding encoder feature maps via skip connections. The decoder outputs a single-channel volume with the same spatial dimensions as the input.

Our model consists of 13, 364, 689 parameters in total and visual schematic of our method is presented in Figure 1.

**Figure 1.**
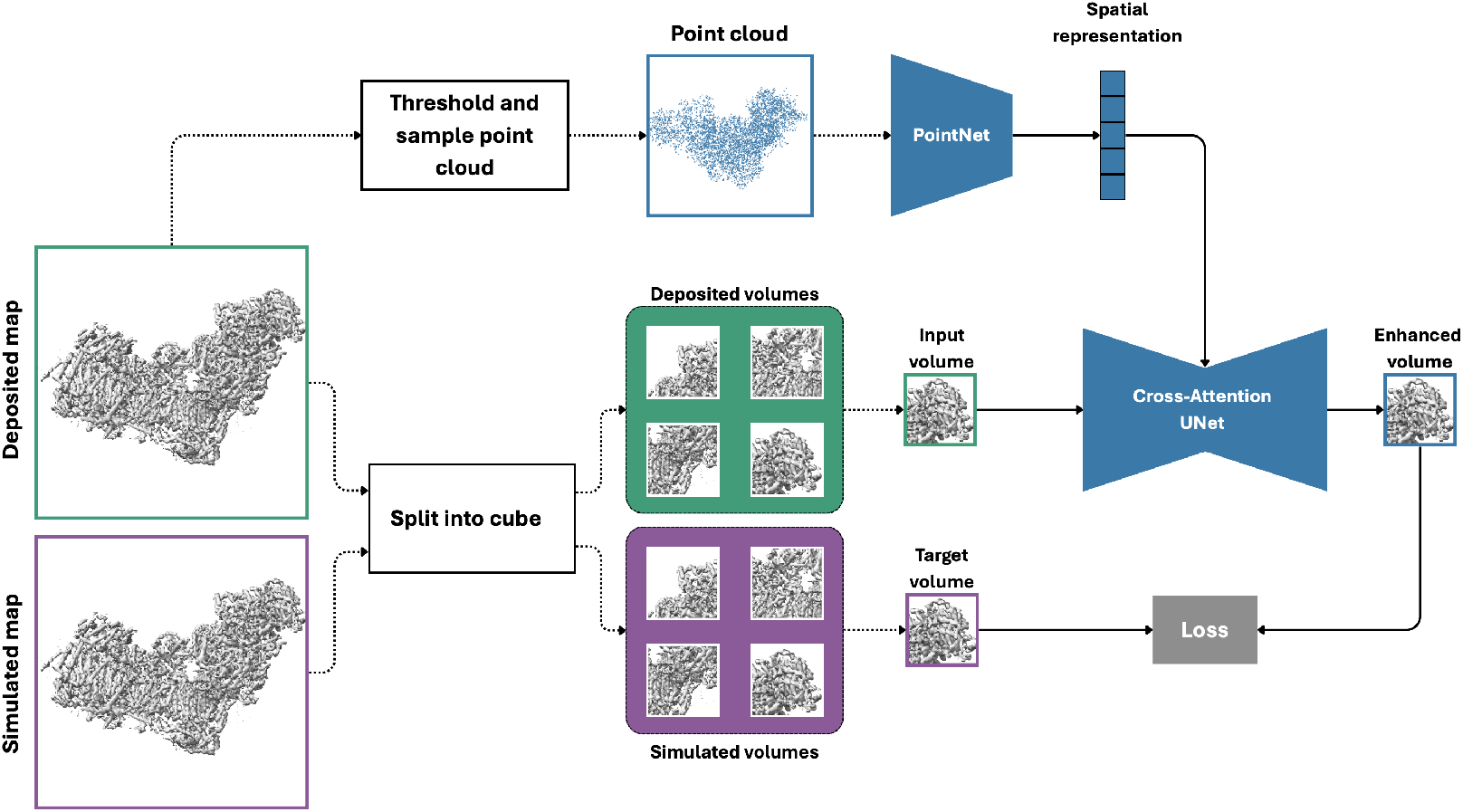
Graphical schema of CryoPC. The deposited map and its simulated counterpart are first divided into corresponding volume cubes. The deposited map is thresholded, and a point-cloud representation is sampled. Each input cube is processed by a UNet, which produces an enhanced volume that is compared with its simulated target. In parallel, a spatial representation extracted by a PointNet is injected into the UNet through a cross-attention mechanism.

**Figure 2.**
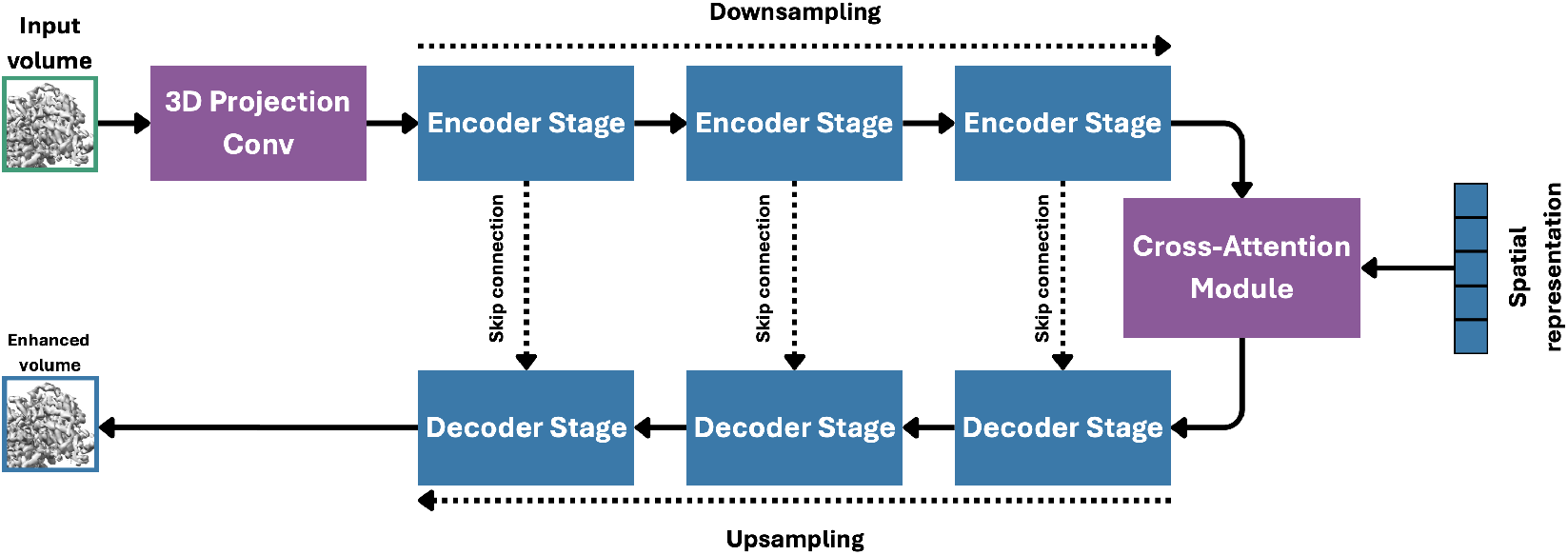
An architecture of the Cross-Attention UNet used in CryoPC. The diagram illustrates the encoder-decoder structure, residual blocks, and the injection of the global point-cloud context via the cross-attention module.

### Experimental setup

We trained three variants of our model: a plain UNet that served as the baseline, our method conditioned on the point-cloud representation, and an extended version of our method that additionally incorporated an SSIM-based regularization component (Wang et al., 2004; He et al., 2023).

Both the UNet baseline and our conditioned model were optimized using the Smooth L1 loss:

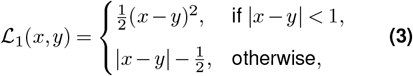

where *x* denotes the predicted volume and *y* the target volume.

For the variant that includes structural regularization, we introduced an SSIM-inspired term:

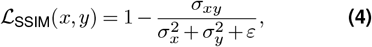

where *σ*_*xy*_ is the covariance between *x* and *y*, 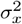 and 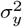 are their respective variances, and *ε* is a small numerical stabilization constant (10^*−*6^ in our experiments). All models were trained on three NVIDIA A100 GPUs with 80 GB of memory each. We employed early stopping based on the validation mean-squared error, terminating training when the validation score did not improve for ten consecutive epochs. Optimization was performed with AdamW (Loshchilov and Hutter, 2019) using a learning rate of 5·10^*−*4^. The learning rate was reduced by a factor of two whenever the validation loss plateaued for four epochs. As a form of data augmentation, input volumes were randomly rotated by 90 degrees. For comparison with previously published methods, we relied on the official implementations provided by the respective authors.

### Evaluation criteria

We used established map–model validation metrics (Afonine et al., 2018; Lawson et al., 2021; Pintilie et al., 2020) to quantify how well each atomic model agrees with its cryo-EM density map and to identify possible inaccuracies or ambiguities in the enhanced map. The Metrics were first computed for deposited maps (post-processed) relative to their reference pdb model and then recalculated for enhanced maps using the same pdb models. Comparing these results allowed us to assess how faithfully each map enhancement correlates to reference pdb model and to characterize the structural improvements introduced by the enhancement.. The metrics used are summarized below. The map–model Fourier shell correlation (FSC) was calculated with `phenix.mtriage` (Afonine et al., 2018). Here, the experimental map is compared in Fourier space with a model-derived map using the FSC at 0.143 resolution (deposited), yielding a curve that reports model–map agreement as a function of spatial frequency. The map–model FSC (FSC_model_) value at 0.5 is correlating to the gold-standard half-map FSC value at 0.143 (Chen et al., 2013; Rosenthal and Rubinstein, 2015). Therefore, FSC_model_ reflects the agreement between the PDB model and the map and is sensitive to enhancing method accuracy and refinement quality. We report FSC_model_ resolutions at 0.143 and 0.5 thresholds using both masked and unmasked maps, as implemented in `phenix.mtriage`.

As a complementary real-space metric, we used the d_model_ resolution from `phenix`, obtained by correlating model-derived maps at different resolutions (adjusted by a global B-factor) with the experimental map. The resolution giving the highest correlation is reported as d_model_ (Afonine et al., 2018) and reflects global real-space agreement between model and cryo-EM density map.

Because d_model_ is relatively insensitive to local detail, we also used the Q-score to assess local atomic resolvability. Q-scores measure how closely the density around each atom resembles well-resolved Gaussian-like density at the same resolution, with values from −1 to 1 (Pintilie et al., 2020).

The Average Q-score correlate with map resolution, allowing comparison with an expected value derived from an empirical regression:

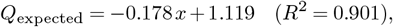

where *x* is a nominal resolution in Å.

To evaluate changes introduced by map enhancement directly from the maps, we used the single-map metric D99 from `phenix`. D99 estimates the highest spatial frequency that meaningfully contributes to the map by identifying the resolution cutoff at which removing higher-resolution Fourier shells reduces self-correlation to 0.99, providing an effective Fourier-based resolution estimate (Afonine et al., 2018).

Finally, real-space interpretability was assessed using map–model cross-correlation scores CC-box, CC-mask, and CC-peaks, computed with `phenix.map_-model_cc` (Afonine et al., 2018). CC-box measures global agreement of the map using the full box, whereas CC-mask restricts the calculation to a model-based masked region (Jiang and Brünger, 1994). CC-peaks compares the strongest density features in experimental and model-derived maps. Together, these metrics provide a complementary evaluation of model agreement and map quality. Following previous studies (Selvaraj et al., 2025; Zhang et al., 2025), we use FSC_model_ at 0.143, FSC_model_ at 0.5, CC-mask, CC-peaks, CC-box, the average Q-score, and the expected Q-score as primary metrics to assess model quality, and treat d_model_ and D99 as complementary metrics for evaluating the quality of individual maps.

## Results

We evaluated all models on an independent test set. During inference, each density map was partitioned into overlapping cubes of size 48 ×48 ×48, processed by the model, and subsequently assembled together to reconstruct the full enhanced map. For constructing point cloud representations, we used the same strategy as for the training. The quality of the resulting enhanced maps was then assessed using the evaluation metrics described in the previous section. Similarly to related methods, we used EMDB deposited maps as a baseline.

### Examining impact of point-cloud conditioning on the map enhancement

Table 1 quantitatively compares deposited cryo-EM maps with three variants of our method: CryoPC trained without point-cloud conditioning, CryoPC with conditioning, and CryoPC with conditioning and additional SSIM regularization, averaged over 120 maps. All trained models substantially improve map quality relative to the deposited maps, as reflected by consistently lower unmasked FSC_model_ values at both the 0.143 and 0.5 thresholds, indicating sharper and more self-consistent maps. Incorporating point-cloud conditioning further reduces FSC_model_ compared to the unconditioned variant, demonstrating that explicit structural guidance provides benefits beyond standard convolutional processing. The addition of SSIM regularization yields the strongest performance across both thresholds. Real-space correlation metrics are also improved for all model variants. In particular, large gains are observed for CC-peaks and CC-box, indicating improved local feature fidelity and global consistency, while CC-mask shows smaller but consistent improvements. Across these metrics, point-cloud–conditioned models outperform the unconditioned CryoPC variant. Q-scores are largely comparable across models, with the SSIM-regularized model achieving the highest average and expected Q-scores, suggesting improved atom-level map interpretability. Overall, these results demonstrate that point-cloud conditioning provides complementary structural guidance for cryo-EM map enhancement, and that SSIM regularization further refines map quality. In the remainder of this paper, we refer to CryoPC + SSIM as **CryoPC**.

**Table 1.** Quantitative comparison of our proposed point cloud models with deposited *primary maps* showing the average of 120 maps for the metrics. Results are shown for the baseline U-Net, CryoPC, and CryoPC trained with SSIM regularization. Both model-map FSC_model_ at 0.143 and at 0.5 are **unmasked** calculated using `phenix.mtriage`.

| | $FSC_{\text{model}}@0.143 \downarrow$ | $FSC_{\text{model}}@0.5 \downarrow$ | CC-mask ↑ | CC-peaks ↑ | CC-box ↑ | Average Q-score ↑ | Expected Q-score ↑ |
| --- | --- | --- | --- | --- | --- | --- | --- |
| Deposited | 3.536 | 5.912 | 0.785 | 0.640 | 0.723 | 0.534 | 0.496 |
| CryoPC w/o Cond. | 2.436 | 3.713 | 0.790 | 0.755 | 0.860 | 0.536 | 0.476 |
| CryoPC | 2.388 | 3.685 | 0.793 | 0.767 | 0.866 | 0.534 | 0.479 |
| CryoPC + SSIM | 2.241 | 3.588 | 0.793 | 0.765 | 0.861 | 0.548 | 0.497 |

### A single-map framework for cryo-EM map enhancement

As cryo-EM map enhancement tools become more widely available, a robust and reproducible framework is needed to assess whether a given method yields meaningful improvements in map quality and interpretability or instead introduces artifacts. To fill this gap, we present a general evaluation framework for individual cryo-EM maps and their enhancements. This framework uses a single refined cryo-EM map and its corresponding atomic model to enable consistent comparisons across methods. An atomic model is often available from previous structural determinations and, or from predictive approaches such AlphaFold (Jumper et al., 2021) or computational methods such as Model-Angelo (Jamali et al., 2023). Additionally, the atomic model must be optimally placed in the experimental cryo-EM map using real-space refinement in `phenix` (Afonine et al., 2018). A set of validation metrics are then computed using the refined map and atomic model with `phenix.mtriage`. These seven metrics are FSC_model_, D99, Q-score, d_model_, and the realspace cross-correlations CC Mask, CC Box, and CC Peaks.

Next, the cryo-EM map is enhanced, but the atomic model is held fixed and not refined against the enhanced map, after which the same metrics are recomputed. This strategy isolates the effect of map enhancement and avoids bias arising from additional model refinement. Because enhancement methods such as CryoPC are trained exclusively on full-map data, applying them independently to half-maps risks introducing correlated noise artifacts and would invalidate the FSC gold-standard (Palovcak et al., 2020; Agarwal et al., 2024). Among the seven metrics, D99 is unique in being entirely model-independent, providing an estimate of effective map resolution by identifying the 1 % of high-frequency Fourier components that do not contribute meaningfully to map information, corresponding to 99 % map cross-correlation. To assess whether enhancement introduces unsupported high-frequency signal, we analyse the single-map D99 metric, which estimates effective map resolution based solely on Fourier content. Empirical analysis shows that moderate sharpening improves D99 for maps at 3.3 Å or lower resolution, whereas over-sharpening beyond 1.9 Å leads to D99 degradation. Across 120 cryo-EM maps spanning nominal resolutions of 2–7 Å, D99 values show strong correlation between deposited and enhanced maps, with only a few outliers, indicating minimal introduction of unsupported high-frequency information.

Of the remaining six metrics, all focus on map-model correlation, but only FSC_model_ evaluates this correlation in Fourier space. Considering FSC_model_ at 0.5 together with D99 provides a complementary assessment of map-model agreement in Fourier space and effective map resolution. To systematically compare enhancement methods, we evaluate metric changes relative to the original map and distinguish between primary and secondary measures based on their physical interpretability. Primary metrics include Fourier- and real-space quantities directly linked to resolution and information content. For example, ΔFSC_model_ at 0.5 and Δ*D*_99_ capture changes in map-model agreement and single-map information content, while ΔQ-score and Δd_model_ quantify local atomic resolvability and global map-model consistency, respectively. Real-space cross-correlation measures (ΔCC-box, ΔCC-mask, and ΔCC-peaks) are treated as secondary metrics. Although these correlations can be computed without explicit resolution estimates, their interpretation is resolution-dependent and sensitive to masking and thresholding choices. Accordingly, CC metrics are most informative for relative comparisons of the same map and serve to support, rather than define, enhancement quality.

Therefore, for an optimal enhancement, the following conditions are assumed to hold for the primary metrics:

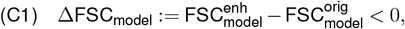

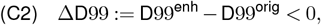

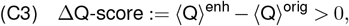

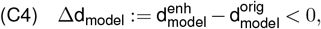

FSC_model_ being taken at 0.5. Additionally, secondary cross-correlation metrics are expected to increase:

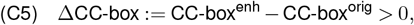

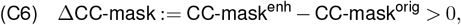

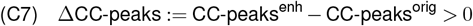

Based on these criteria, the cryo-EM maps are classified as **significantly enhanced** if all primary conditions and all secondary conditions are satisfied, **moderately enhanced** if all primary metrics and at least one secondary metric are satisfied, and **partially enhanced** if one or more primary metrics are not satisfied.

The evaluation of these criteria across the 120 enhanced cryo-EM maps is shown in the scatter plot in Figure 3, where ΔD99 is plotted against ΔFSC_model_ at 0.5. Negative values in both dimensions indicate improved effective resolution and improved map–model agreement, respectively. Accordingly, enhanced maps with ΔD99 *<* 0 exhibit a reduction of unsupported high-frequency Fourier components relative to the deposited maps, consistent with meaningful enhancement of the underlying signal. In contrast, positive ΔD99 values indicate an increase in high-frequency content without corresponding support, suggestive of over-sharpening. Across the dataset, 91 of the 120 enhanced maps show ΔD99 *<* 0, indicating that the majority of enhancements lead to a net gain in meaningful high-frequency information. Only a small number of cases exhibit ΔFSC_model_@0.5 *>* 0, with just two enhanced maps showing a degradation in map–model agreement according to this metric. The overall clustering of points in the lower-left quadrant highlights consistent improvement across complementary Fourier-space measures of effective resolution and map–model agreement.

**Figure 3.**
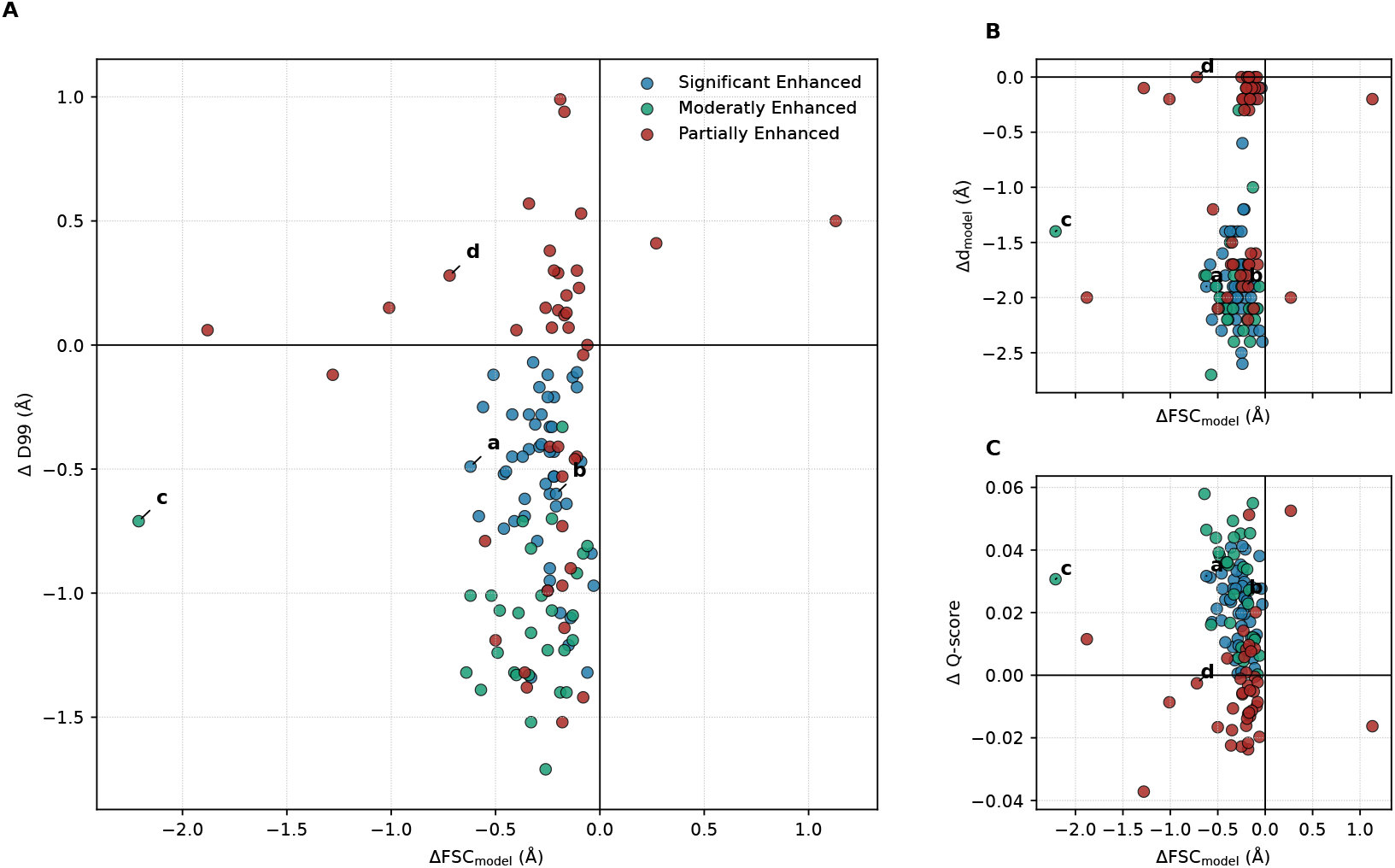
Relationship between resolution-related metrics and map-model agreement for CryoPC-enhanced maps relative to the corresponding deposited primary maps. Metric differences (Δ) are computed as the difference between values obtained for the enhanced maps and those for the original maps. Panel (A) shows the relationship between ΔD99 and ΔFSC_model_ at 0.5, panel (B) shows the relationship between Δ*d*_model_ and ΔFSC_model_ at 0.5, and panel (C) shows the relationship between Δ*Q*-score and ΔFSC_model_ at 0.5. Based on the defined criteria, samples are classified as significantly enhanced, moderately enhanced, or partially enhanced and the maps shown in Figure 4 are labeled a-d.

In general our guidelines provide the user of enhancement tools a method to understand which tools results in the best overall enhancement for their specific cryo-EM map. To highlight our evaluation approach, we are looking at four enhanced maps from the three different regimes shown in Figure 4.

**Figure 4.**
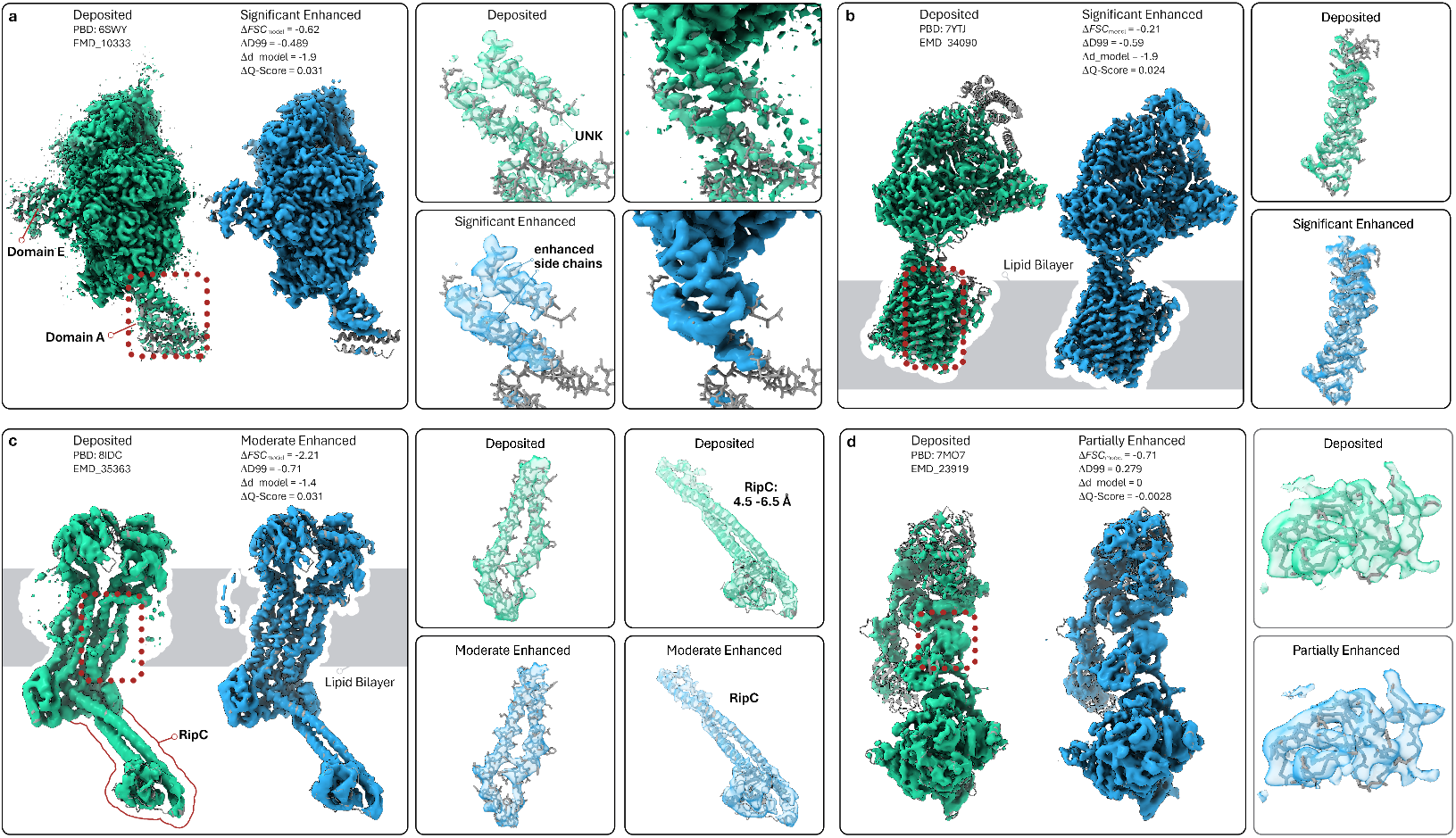
Effects of cryo-EM map enhancement on structural interpretability. Deposited cryo-EM maps are shown in white, enhanced maps in blue, and atomic models as sticks. (a) The GID E3 ubiquitin ligase complex (PDB: 6SWY, EMD-10333) is shown at a resolution of 3.4 Å and compared to a significantly enhanced map. Zoomed views highlight Domain A, where side-chain density is largely absent in the deposited map but becomes clearly resolved after enhancement. The improved density enables tracing of the backbone across most previously unmodelled regions and indicates that approximately 50 of the remaining 106 residues in Domain A can be reliably incorporated into the model. (b) The signal-activated vacuolar transporter chaperone (VTC) complex (PDB: 7YTJ, EMD-34090) is shown at a resolution of 3.0 Å and exhibits significant enhancement. Notably, the transmembrane region embedded in the lipid bilayer shows improved side-chain definition, enabling more reliable model interpretation. (c) The cryo-EM map of the Mycobacterium tuberculosis FtsEX–RipC complex (PDB: 8IPC, EMD-35363) is shown at an overall resolution of 3.9 Å, with local resolution in the RipC domain ranging from 4.5 to 6.5 Å. This map is classified as moderately enhanced. While side-chain features of α-helical residues in the transmembrane region become more pronounced after enhancement, density corresponding to the RipC domain appears smoother, reflecting the limited effect of enhancement at lower local resolution. (d) The cryo-EM map of the 2:2 c-MET/HGF holo-complex (PDB: 7MO7, EMD-23919) is shown at a resolution of 4.8 Å. In contrast to the previous examples, the partially enhanced map exhibits a loss of high-frequency detail relative to the deposited map, with noticeable smoothing and reduced side-chain definition. This example illustrates that enhancement does not universally improve interpretability and may attenuate side-chain information in some cases

### Visual inspection of CryoPC

-enhanced maps To illustrate how CryoPC enhancement occurs in different structural regimes, we analysed several representative cryoEM maps that were classified by our framework as being significantly, moderately or partially enhanced. First, we examined the structure of the active GID E3 ubiquitin ligase complex (minus Gid2 and the Gid9 RING domain) from Saccharomyces cerevisiae (Figure 4a), which our criteria classify as significantly enhanced. In chain E, which corresponds to FYV10 of the GID complex 48 residues are not identified in the sequence (annotated as residues tpye UNK) and are modeled with only the peptide backbone and c-alpha atoms. Overlaying this with the deposited cryo-EM density (green) and the CryoPC-enhanced map (blue) clearly reveals a continuous Cα trace for at least 42 of those residues. Visual inspection suggests that approximately 40 side chains are at least partially interpretable in the enhanced density. Similarly, in chain A, which corresponds to the vacuolar import and degradation protein A (Vid30), of which 106 residues contain only the backbone atoms (no side chains) unmodeled in the deposited structure. Of these, roughly 50 residues show discernible side chain features in the enhanced map.

Next, we inspected the vacuolar transporter chaper-one (VTC) complex, which is a membrane protein from Saccharomyces cerevisiae containing five transmembrane helices (Figure 4b). This map is also classified as significantly enhanced. Comparing the deposited map (green) with the enhanced map (blue) reveals a clear improvement in structural detail within the trans-membrane region. In particular, side-chain features along α-helical segments normally embedded in detergent or lipid environments become markedly more distinct upon enhancement. More generally, across multiple membrane-protein datasets, we observe that \our{enhancement particularly benefits α-helical transmembrane regions, where high-resolution features would otherwise be attenuated by surrounding solvent or detergent density.

To illustrate a moderately enhanced dataset, we analysed the cryo-EM structure of the Mycobacterium tuberculosis FtsEX complex (Figure 4c), which is a non-canonical regulator with high basal ATPase activity. The enhanced map reveals a sharpening effect in the transmembrane helices compared to the deposited map, which is similar to that observed for the VTC complex. However, in the RipC domain, side-chain features appear to be improved less consistently, and in some regions they are diminished. This behaviour at the RipC domain is likely attributable to the low local resolution of 4.5 Å to 6.5 Å with respect to the resolution of the remaining cryoEM map (3.9 Å), where the trans-membrane region holds the highest resolution. In the resolution regime of the RipC domain the side-chain information is inherently weak and difficult to interpret.

Lastly, Figure 4d shows the cryo-EM map of the c-MET receptor, a receptor tyrosine kinase classified as partially enhanced. The deposited map exhibits local resolution estimates ranging from 4.5 Å to 6.5 Å. The enhanced map reveals a smoothing of side-chain features that were already weakly defined in the deposited reconstruction. This suggests that, when the underlying signal is insufficient, CryoPC does not artificially introduce high-frequency features. Rather, these results demonstrate that CryoPC performs optimally at map resolutions where side-chain information is present. For significantly and moderately enhanced cryo-EM maps, we observed a mean resolution of approximately 3.3 Å, with a total of 77 maps showing a resolution better than 4 Å. Of these, only three cryoEM maps had a resolution between 4 and 4.5 Å. For partially enhanced maps, 27 had a resolution better than 4 Å, and 15 had a resolution larger than 4 Å, resulting in an average resolution of 3.9 Å.

### Comparison with existing deep learning models

We evaluated CryoPC against four established methods: DeepEMEnhancer (Sánchez-García et al., 2020), EMReady (He et al., 2023), CryoTEN (Selvaraj et al., 2025), and CryoSAMU (Zhang et al., 2025) using our test-set of 120 maps. To ensure consistency with recent standards, we adopted the quantitative results reported by Selvaraj et al. (2025) for DeepEMEnhancer, EMReady, and CryoTEN, while ensuring all comparisons were based on the same set of evaluation metrics for the original deposited maps. Additionally, we assessed scalability and robustness using a subset of six randomly selected maps (ranging from 10^6^ to 10^8^ voxels), with processing times averaged over 10 independent runs. Current evaluation protocols typically report average metrics across datasets and highlight selected examples, often focusing on a single quality indicator. While useful for high-level comparisons, this approach obscures protein-specific structural heterogeneities and fails to capture nuances in individual map enhancements. Consequently, while average metrics provide a necessary general baseline, they are insufficient to fully characterize enhancement quality, and a rigorous framework for one-to-one map comparison remains lacking.

To ensure consistent map–model evaluation, each enhanced map was assessed using its corresponding deposited atomic model without additional refinement. As shown in Table 3, CryoPC achieves clear overall improvements in map–model agreement across both Fourier- and real-space metrics, indicating systematic enhancement of cryo-EM densities. In terms of FSC_model_, CryoPC consistently improves upon deposited maps and performs competitively with state-of-the-art methods, achieving substantially lower FSC_model_ 0.5 than DeepEMEnhancer and CryoSAMU, slightly better than CryoTEN, while EMReady being the best. CryoPC also maintains high correlation coefficients across all CC metrics, matching EMReady and exceeding DeepEMEnhancer, CryoTEN and CryoSAMU, indicating robust global and local agreement between atomic models and enhanced maps.

Beyond reconstruction quality, CryoPC demonstrates favorable computational efficiency. As reported in Table 2, CryoPC processes maps significantly faster than EMReady and DeepEMEnhancer, which require several minutes per volume, while remaining on par with the fastest methods, CryoTEN and CryoSAMU. With its strong quantitative performance, robustness across map sizes, and competitive runtime, CryoPC offers an attractive trade-off between accuracy and computational efficiency, making it well suited for large-scale cryo-EM map enhancement workflows.

**Table 2.** Comparison of processing time across different models. The performance was analyzed on a subset of six randomly selected maps ranging in size from 10^6^ to 10^8^ voxels. The result in the table is the average processing time from 10 runs.

|  | DeepEMEnhancer | EMReady | CryoTEN | CryoSAMU | CryoPC (Ours) |
| --- | --- | --- | --- | --- | --- |
| Average Processing Time (s) | 512.46 | 910.27 | 27.37 | 23.09 | 37.94 |

**Table 3.**
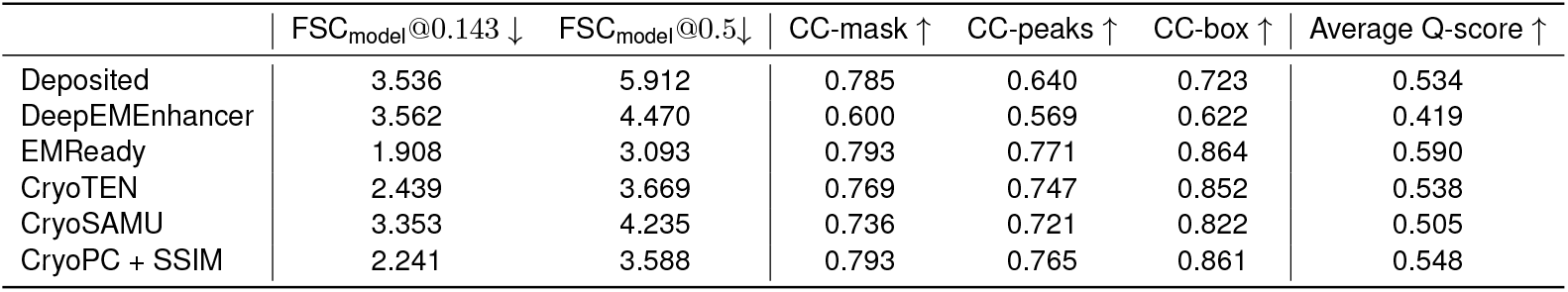
Quantitative comparison of our proposed models with deposited reference maps. Results are shown for the DeepEMEnhancer, EMReady, CryoTEN, CryoSAMU and CryoPC trained with SSIM regularization. Both FSC_model_ at 0.143 and FSC_model_ at 0.5 are **unmasked** FSC_model_ calculated using `phenix.mtriage`.

| | $FSC_{model}@0.143 \downarrow$ | $FSC_{model}@0.5 \downarrow$ | CC-mask ↑ | CC-peaks ↑ | CC-box ↑ | Average Q-score ↑ |
| --- | --- | --- | --- | --- | --- | --- |
| Deposited | 3.536 | 5.912 | 0.785 | 0.640 | 0.723 | 0.534 |
| DeepEMEnhancer | 3.562 | 4.470 | 0.600 | 0.569 | 0.622 | 0.419 |
| EMReady | 1.908 | 3.093 | 0.793 | 0.771 | 0.864 | 0.590 |
| CryoTEN | 2.439 | 3.669 | 0.769 | 0.747 | 0.852 | 0.538 |
| CryoSAMU | 3.353 | 4.235 | 0.736 | 0.721 | 0.822 | 0.505 |
| CryoPC + SSIM | 2.241 | 3.588 | 0.793 | 0.765 | 0.861 | 0.548 |

## Conclusions

In this paper, we introduced CryoPC, a novel neural network architecture conditioned on point-cloud representations designed to enhance cryo-EM maps. Our experiments demonstrate that integrating point-cloud data improves the model’s enhancemewnt capabilities. Crucially, CryoPC achieves a favorable balance between computational efficiency and output fidelity, proving to be an effective tool for map enhancement. Furthermore, we presented a comprehensive evaluation framework to rigorously assess the quality of cryo-EM maps. In future work, we plan to continue exploring additional conditioning mechanisms and other neural network architectures to further refine our approach.

## Author contributions statement

Author Contributions K.K. conceived the study and designed the experiments. K.K. and M.C. conducted the experiments and performed data collection. A.R. and M.Z. provided the essential resources and computational infrastructure. K.K. and L.F. performed the formal analysis and prepared the visualizations. All authors contributed to the writing, review, and final editing of the manuscript.

## Acknowledgments

The authors would like to express their sincere gratitude to Joel Selvaraj for his valuable correspondence and assistance with the CryoTEN code and to Jason Greenwald for his insightful input on the manuscript.

## Data availablity

The code as well as the model’s weights are available on https://github.com/konrad-karanowski/cryopc-enhancement.

